# GPT-4V shows human-like social perceptual capabilities at phenomenological and neural levels

**DOI:** 10.1101/2024.08.20.608741

**Authors:** Severi Santavirta, Yuhang Wu, Lauri Suominen, Lauri Nummenmaa

## Abstract

Humans navigate the social world by rapidly perceiving social features from other people and their interaction. Recently, large-language models (LLMs) have achieved high-level visual capabilities for detailed object and scene content recognition and description. This raises the question whether LLMs can infer complex social information from images and videos, and whether the high-dimensional structure of the feature annotations aligns with that of humans. We collected evaluations for 138 social features from GPT-4V for images (N=468) and videos (N=234) that are derived from social movie scenes. These evaluations were compared with human evaluations (N=2254). The comparisons established that GPT-4V can achieve human-like capabilities at annotating individual social features. The GPT-4V social feature annotations also express similar structural representation compared to the human social perceptual structure (i.e., similar correlation matrix over all social feature annotations). Finally, we modeled hemodynamic responses (N=97) to viewing socioemotional movie clips with feature annotations by human observers and GPT-4V. These results demonstrated that GPT-4V based stimulus models can also reveal the social perceptual network in the human brain highly similar to the stimulus models based on human annotations. These human-like annotation capabilities of LLMs could have wide range of real-life applications ranging from health care to business and would open exciting new avenues for phycological and neuroscientific research.

## Introduction

Humans encounter complex social situations every day and fast perception of the social context is vital when navigating in the social world. Rapid social perception is based on eight main social dimensions allowing quick inference and reacting to changes in the dynamic situation (Santavirta et al., 2024). Rapid extraction of social information is necessary for understanding others’ intentions, predicting their behavior and adapting dynamically to the social environment. Large-language models (LLM) have developed rapidly in recent years for chat-based interaction with humans and they have shown promising capabilities across disciplines including social cognition. LLMs excel in complex knowledge tests such as Uniform Bar Examination (Katz et al., 2024), the United States Medical Licensing Exam (Kung et al., 2023) and also general IQ tests (King, 2023). Currently, OpenAI’s GPT-4 model is one of the most widely used and researched model that is capable of achieving high and near human performance across disciplines (Bubeck et al., 2023) and has outperformed other state-of-the-art models in many psychological tasks (Almeida et al., 2024). Recent studies have explored how LLMs can be used for advancing behavioral science (Demszky et al., 2023). With the increase of the “humanness” of the models, psychological research has been starting to explore whether AI models can even replace human participants (Dillion et al., 2023; Horton, 2023; Sohail & Zhang, 2025). Studies have found that LLMs can infer emotions from text input (Huang et al., 2023; Tak & Gratch, 2024; X. Wang et al., 2023), simulate collective intelligence of multiple humans (Aher et al., 2023), simulate opinions of different sociodemographic groups (Argyle et al., 2023), make human-like economical choices (Horton, 2023), reach human-like performance in many cognitive tasks (Binz & Schulz, 2023), emulate personality (Y. Wang et al., 2025) and simulate human-like theory of mind (Strachan et al., 2024).

However, a large bulk of human social cognition is based on audiovisual input from the environment, rather than written text. Recently, LLMs have been coupled with advanced vision and speech-to-text models which enable them to describe the contents of images and videos. Popular models with vision capabilities are currently OpenAI’s GPT-4V (Yang et al., 2023) and Google’s Gemini 2.5 (Gemini Team et al., 2024) among others. In this study, we question whether LLMs are already capable of inferring complex social characteristics (such as dynamics of social interaction) that are based on contextual and indirect inference from the scene. Such human-like high-level social perception from videos and images could have significant applications. AI could, for example, monitor patients’ physical and psychological well-being in healthcare. In customer service, AI could replace the common customer satisfaction inquiries and instead extract this information from the interaction between customers and their services or products. In turn, AI-based video surveillance could identify conflicts and anticipate them even before they occur.

For research purposes the scalability of the LLMs would allow unprecedented efficiency in data collection that could be used for generating large-dimensional stimulus models for new and retrospective datasets. It is suggested that LLMs could transform the computational social science with its scalable ability to annotate any kind of textual input, opening the current bottleneck of data annotations of large datasets (Ziems et al., 2023). Functional brain imaging has previously mapped the high-dimensional brain representations for multiple cognitive and social phenomena (Huth et al., 2016, 2012; Koide-Majima et al., 2020; Lettieri et al., 2019; Saarimäki et al., 2025; Santavirta et al., 2023; Tarhan & Konkle, 2020). Such large-scale analyses require pooling massive amounts of human evaluations from the stimulus contents for setting up the stimulation models for functional imaging experiments. Stimulus annotation is labor intensive especially when high temporal resolution (e.g. video stimuli) is needed. For example, high-dimensional annotations for mapping the representational space for social perception in images and videos used in this study required approximately 1100 hours of labor from the volunteers (Santavirta et al., 2024). Augmenting human ratings with automated image or video evaluations would massively scale up the possibilities to map high-dimensional representations in the brain. Importantly, this would also allow reanalysis of many already acquired functional imaging datasets which however lack detailed annotation of the stimuli and thus detailed high-dimensional stimulus models.

Here we show that GPT-4V can reproduce concrete and complex human social perceptual annotations. We also show their practical usefulness for neuroimaging as these annotations can be used as reliable stimulus models for modeling neural basis of social perception in functional magnetic resonance imaging data. We asked GPT-4V to provide evaluations for 138 social perceptual features from 468 static images and 234 short video scenes depicting people in different social situations. We compared how similarly GPT-4V evaluated the features compared to the reference human evaluations pooled from 2254 human observers. We also demonstrate that the agreement between GPT-4V and humans is reflected in the neural level in a large fMRI dataset of 97 subjects watching short videos with social content (96 videos) by comparing the neural response profiles based on stimulus models derived from GPT-4V versus human observers. The results show high consistency in social feature annotation between GPT-4V and humans supporting the advanced capabilities of GPT-4V in extracting complex social information from audiovisual input.

## Methods

The stimulus, evaluated social features, human data collection procedure, human sample size estimation and data quality control are described in detail in the previous publication of the social perceptual taxonomy in humans (Santavirta et al., 2024). **Figure 1** shows the analytical workflow of the study.

**Figure 1.**
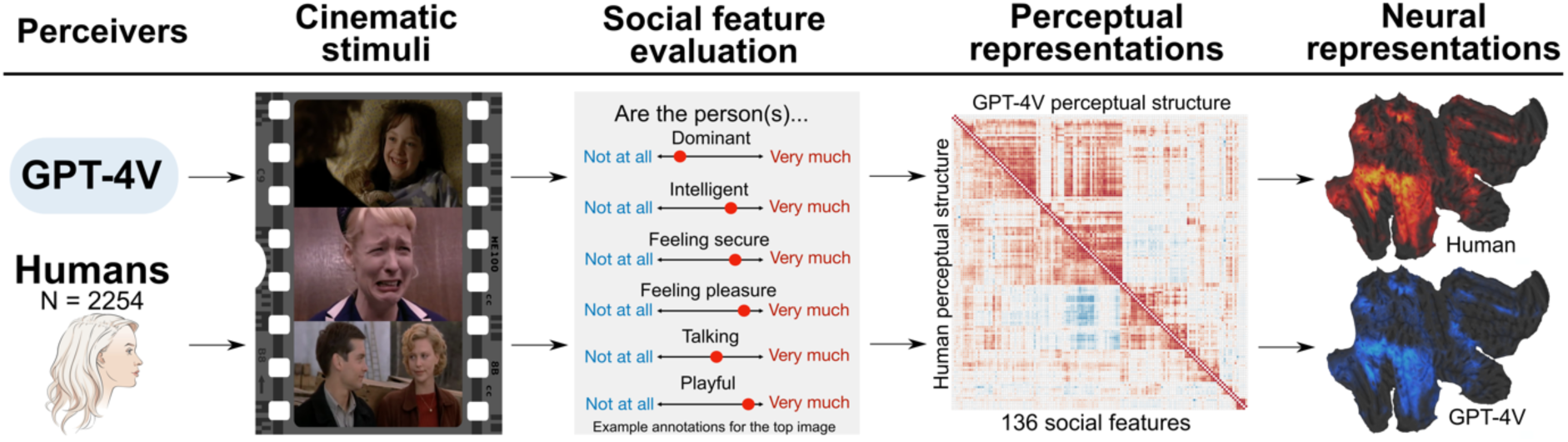
Analytical workflow of the study. GPT-4V and humans evaluated the presence of 138 social features from images and movie clips and the similarity of the evaluations between GPT-4V and humans was investigated. We then tested the similarity of the perceptual ratings and the perceptual structure between humans and GPT-4V (136 features were analyzed since human did not perceive two features in the stimuli at all.). These evaluations were then used to generate stimulus models for mapping neural representations of social perception in a large fMRI dataset (n=97) and to compare the resulting mappings between humans and GPT-4V.

### Stimulus

To estimate whether GPT-4V can retrieve social features from videos and images similarly as humans, we tested its performance in large datasets of images and videos containing socioemotional scenes. Movies are effective for studying social perception in controlled settings since movies typically present natural, socially important episodes with high affective intensity and frequency. Using movies as stimuli, social neuroscience has successfully studied neural representations of social and emotional processing (Adolphs et al., 2016; Lahnakoski et al., 2012; Nummenmaa et al., 2023; Saarimäki, 2021; Santavirta et al., 2023). Hence, we selected short movie clips primarily from mainstream Hollywood movies (N=234) as the video stimulus. The same dataset was used previously for defining the basic eight taxonomy for social perception (Santavirta et al., 2024). The average duration of the movie clips was 10.5 seconds (range: 4.1 - 27.9 sec) with a total duration of 41 minutes. The image dataset was generated by capturing two clear frames from each of the movie clips. A research assistant not familiar with the study was instructed to select two clear frames, preferentially from two different scenes if the clip contained more than one shot. The image stimulus set contained a total of 468 images. See **Table SI-1** for the descriptions of the movie clips.

### Evaluated social features

The initial feature set contained a total of 138 social features that were selected to cover a broad range of perceptual properties relevant for social interaction and to identify whether GPT-4V can evaluate the presence of some social features in the movie clips more consistently with humans than others. This feature set has been previously validated to broadly represent the human social perceptual space (Santavirta et al., 2024). Features based on previous theories of social cognition were selected from the following broad perceptual categories: person’s personality traits, person’s physical characteristics, person’s internal situational states, somatic functions, sensory states, qualities of the social interaction, communicative signals and persons’ movement as the broad perceptual categories. See **Table SI-2** for the full list of evaluated features.

### Large language model

GPT-4V was selected as the tested model family since they are currently the most popular and researched models. Their API is easy to use and affordable. Initially the data were collected with older models (images: gpt-4-1106-vision-preview, videos: GPT-4-turbo-2024-04-09), but the presented results are updated to the newest vision model (gpt-4.1-2025-04-14). A model comparison of the results can be found in **Table SI-3**.

### Human perceptual reference evaluations

For each image and video, we collected 10 independent perceptual ratings for each social feature from human observers. Participants were instructed to evaluate the prominence of each feature on an abstract scale between end points “Not at all” and “Very much” using a continuous slider. To match the evaluations with the numerical evaluations of GPT-4V, the abstract slider responses were transformed to numerical values between 0 and 100. To minimize cognitive load and participant exertion, a single participant only evaluated 6 - 8 features from a subset of the stimuli (39 videos or 78 images) which took approximately 30 minutes of each participant’s time. The order of the images and videos was randomized for each participant to ensure that the evaluation order does not bias the population level results. The dataset with images (i.e. movie frames) included 1094 participants from 56 nationalities with approximately 654 000 data points. 448 participants were female (41 %) and the median age of the participants was 32 years (range: 18 - 77 years). The reported ethnicities of the participants were: White (770, 70.4 %), Black (194, 17.7 %), Mixed (55, 5.0 %), Asian (51, 4.7 %) and Other (20, 1.8 %). The dataset with video clip evaluations included 1 096 participants from 60 nationalities with approximately 327 000 data points. 515 participants were female (46, 9 %) and the median age of the participants was 28 years (range: 18 - 78 years). The reported ethnicities were: White (788, 71.9 %), Black (136, 12.4 %), Mixed (81, 7.4%), Asian (47, 4.3 %) and Other (30, 2.7 %). Based on the human evaluations, the stimulus did not contain “Coughing/sneezing” and “Vomiting/urinating/defecating” at all. These features were excluded from the analyses and the results are based on the comparison of perception in 136 social perceptual features. The reference data and its acquisition has been reported in detail previously (Santavirta et al., 2024).

### GPT-4V image perception experiment

We designed the input prompts for GPT-4V to follow the human instructions as closely as possible, while assuring that the prompts were accessible by GPT-4V (see section “Defining the social feature evaluation instructions (prompt) for GPT-4V” in the Supplemental materials). The main difference compared to humans in the data collection was that GPT-4V was asked to output numerical ratings between 0 and 100 instead of using the abstract slider that humans used and GPT-4V evaluated all features at once from the given image (or video in the following video experiment). GPT-4V API was used to collect evaluations for each image one-by-one. Each image was added to the same prompt alongside with the instructions. While ChatGPT can remember previous conversations within a chat, the API should not have memory over consecutive requests (https://community.openai.com/t/does-the-open-ai-engine-with-gpt-4-model-remember-the-previous-prompt-tokens-and-respond-using-them-again-in-subsequent-requests/578148), which ensures that that GPT-4V rates each frame independently. Sometimes GPT-4V refused to provide ratings mainly for sexual stimuli presumably due to content moderation (Respond: “I’m sorry, but I can’t provide assistance with this request”) while being able to provide ratings with a new enquiry. Therefore, the failed images were fed to the API again until GPT-4V refused to return any new data. The still missing data points were excluded from the data before analyses.

As a stochastic model GPT-4V does not provide the same output for a given prompt every time. Hence, it is suggested that pooling multiple responses, just like recruiting multiple human participants in traditional experiments, can increase the generalizability of the responses (Demszky et al., 2023). Hence, we repeated the data collection procedure to get five full datasets of GPT-4V responses and found that averaging over multiple collection rounds increased the agreement in the ratings, while the agreement increase started to plateau after a few data collections rounds (see section “Collecting several rounds of GPT-4V data increases the agreement between GPT-4V and humans” in the Supplemental materials). After repeated data collection in five collection rounds GPT-4V failed to provide ratings for 17 out of 468 images (3.6 %) for all five datasets, and these images were thus excluded from the further analyses. 12 of the failed images contained sex, two contained blood, one depicted a man who is about to hang himself and two depicted humans in everyday activities. The data were collected in May 2025 using the “gpt-4.1-2025-04-14” model (https://platform.openai.com/docs/models). The average ratings over the five collection rounds were used to compare the social perceptual capabilities of GPT-4V against the human evaluations.

### GPT-4V video perception experiment

At the time of data collection (June 2025), GPT-4V could not process video input directly. Therefore, we extracted eight frames uniformly from video clips. The speech in the movie clips was extracted and transformed to text using the “whisper-1” model (https://platform.openai.com/docs/guides/speech-to-text). For videos without any conversation, the model gave unreliable responses (e.g. “Thanks for watching”) which were manually removed. All transcripts were manually checked and confirmed that the videos with suspicious transcripts indeed did not contain any speech. The final “video perception” prompt for social feature evaluation included the transcript alongside the captured frames. The rating data were collected for each movie clip in separate GPT-4V API requests. Five full datasets of GPT-4V responses were collected and the final dataset was formed as average over the five collection rounds. Averaging increased the agreement between human and GPT-4V responses also in the video perception data (see section “Collecting several rounds of GPT-4V data increases the agreement between GPT-4V and humans” in the Supplemental materials). After repeated data collection in five collection rounds GPT-4V failed to provide ratings for 6 out of 234 videos (2.6 %) for all five datasets, and these videos were thus excluded from further analyses. All these videos contained sex. The data was collected in May 2025 using the updated “gpt-4.1-2025-04-14” model (https://platform.openai.com/docs/models).

### Analysis of the rating consistency between GPT-4V and humans for social perception

First, we analyzed how similarly GPT-4V and humans perceive the social context in naturalistic scenes. Similar analysis protocol was used for analyzing the image and video datasets. To evaluate how similarly GPT-4V perceived social features compared to humans we calculated “the agreement of GPT-4V” -index as the Pearson correlation between GPT-4V evaluations and the human average ratings individually for each social feature. There is considerable variation in the consistency of social perceptual evaluations between different individuals (Santavirta et al., 2024). Some social features are concrete (such as sex) and yield nearly 100% consistency across human observers whereas others are more ambiguous (such as moral righteousness) yielding larger variation. Still, in our neuroimaging dataset and commonly in other neuroimaging experiments, the neuroimaging data of independent participants are modelled with population average annotations derived from another sample of participants. Hence, the suitable benchmark for the agreement of GPT-4V is the “intersubject consistency” -index which was computed as the mean of Pearson correlations between individual observers’ ratings and the population mean of all other human participants. The intersubject consistency thus estimates how consistently single subjects’ perceptual ratings track the population average (i.e., how similarly a participant in the fMRI scanner likely perceives the social world compared to group of others). This was used as the benchmark against which the GPT-4V ratings were compared to. Since the human data contained ten independent annotations for each item, we also calculated a “group-level consistency” of human average ratings by splitting the data into two groups of five annotations and calculating the average rating for these two groups. Then, correlation between the two groups was calculated for all possible combinations of independent groups of five annotations. This group-level agreement was used as a higher benchmark against the agreement of GPT-4V.

### Consistency analysis of the social perceptual structure between GPT-4V and humans

Humans show a generalizable low-dimensional perceptual structure over the analyzed 136 features (Santavirta et al., 2024). Following the dimension reduction methods developed previously (Santavirta et al., 2024), we compared the similarity of the correlation matrices between GPT-4V and human social feature ratings by calculating the Pearson correlation between the GPT-4V and human correlation matrices of social feature ratings. The statistical significance of the correlation was tested with a non-parametric Mantel test with 1 000 000 permutations (Mantel, 1967) using the Ape R package (https://rdrr.io/cran/ape/man/mantel.test.html). Secondly, we used principal coordinate analysis (PCoA) to map the social perceptual structures based on GPT-4V and human ratings separately and then compared how similar is the social perceptual structure of GPT-4V with the human social perceptual structure. PCoA was implemented using the R function cmdscale (https://www.rdocumentation.org/packages/stats/versions/3.6.2/topics/cmdscale) with a Pearson correlation matrix of the individual social features as input. Correlation was selected as the distance metric for PCoA, since it measures the covariance of the ratings between features more accurately than Euclidean distance used in standard principal component analysis (PCA). PCoA transforms the original correlation structure into orthogonal principal components. Each original social feature contributes (loads) to the principal components and based on these loadings it is possible to infer which kind of social information the components contain. By correlating the component loadings between GPT-4V and human derived PCoA components, it is possible to investigate how similar the social perceptual structure is between the results. We hypothesized that the first eight principal components would show consistency between GPT-4V and humans since only the first eight components were identified to explain meaningful variance in the social perceptual structure in human data (Santavirta et al., 2024).

### Neuroimaging experiment

The neuroimaging dataset with 102 volunteers has been reported in detail previously (Santavirta et al., 2023). The exclusion criteria included a history of neurological or psychiatric disorders, alcohol or substance abuse, BMI under 20 or over 30, current use of medication affecting the central nervous system and the standard MRI exclusion criteria. Two additional subjects were scanned but excluded from further analyses because of unusable MRI data due to gradient coil malfunction. Two subjects were excluded because of anatomical abnormalities in structural MRI and additional three subjects were excluded due to visible motion artefacts in preprocessed functional neuroimaging data. This yielded a final sample of 97 subjects (50 females, mean age of 31 years, range 20 – 57 years). To map brain responses to different social features, we used our previously validated socioemotional “localizer” paradigm that allows reliable mapping of various social and emotional functions (Karjalainen et al., 2017; Lahnakoski et al., 2012; Nummenmaa et al., 2021; Santavirta et al., 2023). The experimental design and stimulus selection has been described in detail in the original study with this setup (Lahnakoski et al., 2012). The subjects viewed a medley of 96 movie clips (median duration 11.2 s, range 5.3 – 28.2 s, total duration 19 min 44 s) and 87 of these movie clips were included in the stimulus set for GPT-4V social feature evaluation enabling mapping the neural representations for GPT-4V social perception.

### Ethics statement

All neuroimaging subjects gave an informed, written consent and were compensated for their participation. The ethics board of the Hospital District of Southwest Finland had approved the protocol, and the study was conducted in accordance with the Declaration of Helsinki.

### Neuroimaging data acquisition and preprocessing

MR imaging was conducted at Turku PET Centre. The MRI data were acquired using a Phillips Ingenuity TF PET/MR 3-T whole-body scanner. High-resolution structural images were obtained with a T1-weighted (T1w) sequence (1 mm3 resolution, TR 9.8 ms, TE 4.6 ms, flip angle 7°, 250 mm FOV, 256 × 256 reconstruction matrix). A total of 467 functional volumes were acquired for the experiment with a T2∗-weighted echo-planar imaging sequence sensitive to the blood-oxygen-level-dependent (BOLD) signal contrast (TR 2600 ms, TE 30 ms, 75° flip angle, 240 mm FOV, 80 × 80 reconstruction matrix, 62.5 kHz bandwidth, 3.0 mm slice thickness, 45 interleaved axial slices acquired in ascending order without gaps).

MRI data were preprocessed using fMRIPprep 1.3.0.2 (Esteban et al., 2019). The following preprocessing was performed on the anatomical T1-weighted (T1w) reference image: correction for intensity non-uniformity, skull-stripping, brain surface reconstruction, and spatial normalization to the ICBM 152 Nonlinear Asymmetrical template version 2009c (Fonov et al., 2009) using nonlinear registration with antsRegistration (ANTs 2.2.0) and brain tissue segmentation. The following preprocessing was performed on the functional data: coregistration to the T1w reference, slice-time correction, spatial smoothing with a 6-mm Gaussian kernel, non-aggressive automatic removal of motion artifacts using ICA-AROMA (Pruim et al., 2015), and resampling of the MNI152NLin2009cAsym standard space.

### Modeling the similarity of the neural representations for social perception between GPT-4V and humans

To test whether GPT-4V derived stimulus models produce similar neural representations compared to those based on human observations, we first modeled the hemodynamic responses measured with fMRI separately with GPT-4V and human derived regressors for social features and then compared the similarity of the results. Simple regressions were conducted separately using SPM12 (Wellcome Trust Center for Imaging, London, UK, http://www.fil.ion.ucl.ac.uk/spm). Each social feature was convolved with a canonical double-gamma hemodynamic response function before analyses. Then, social feature regressors were independently fitted to each participant’s voxel-level fMRI data (first level analysis, massive univariate approach). The resulting subject-level β-coefficient-maps were subjected to group-level analysis to identify the population level association between social features and the hemodynamic response. One-sample t-tests were used to statistically threshold the population level results.

The feature-specific similarity in the neural representations was assessed by i) correlating the unthresholded β-coefficient-maps between GPT-4V and humans for testing the overall spatial similarity of the result distributions, and by ii) calculating the positive and negative predictive values (PPV = TP / [TP + FP], NPV = TN / [TN + FN]) of the statistically thresholded GPT-4V results to assess how reliable the thresholded GPT-4V response maps are for different social features. PPVs and NPVs were calculated for the positive main effect of the social features considering the human derived results as the ground truth. They are reported for conservatively threshold results (voxel-level FWE-corrected, p < 0.05) as well as for results with lenient statistical threshold (p < 0.001, uncorrected). To reveal the general social perceptual network, we calculated the social cumulative map as the sum of social features that were positively associated with the hemodynamic response (p < 0.001, uncorrected) in each voxel and compared the cumulative maps between GPT-4V and human derived results.

## Results

The overall similarity of the social feature ratings between GPT-4V and human average is visualized in **Figure 2**. The correlation between GPT-4V and human ratings over all social features was 0.79 for images and also 0.79 for videos indicating high consistency in the ratings.

**Figure 2.**
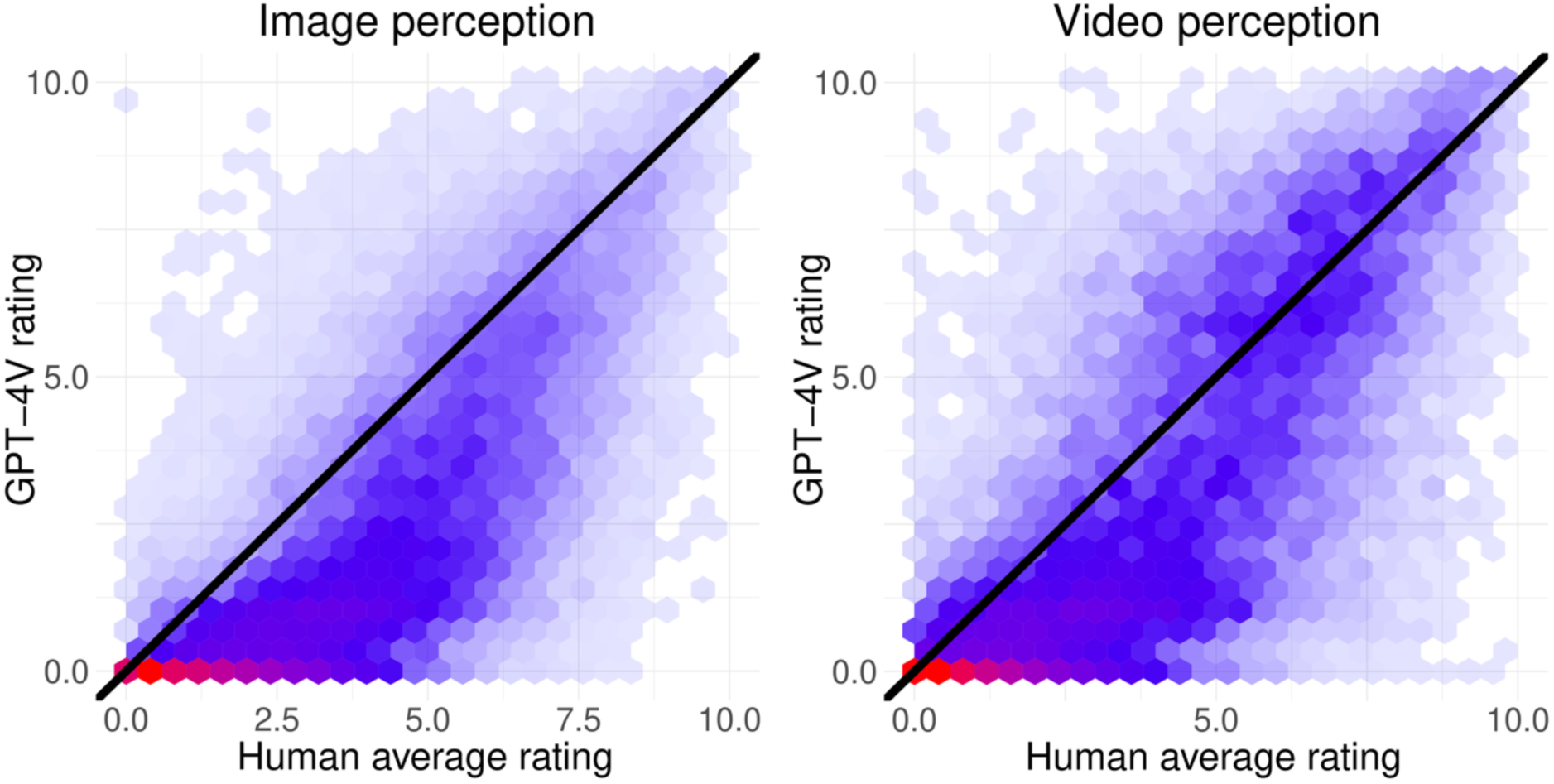
Density plots of GPT-4V ratings against the human average ratings calculated as the average over ten human annotations.

### GPT-4V rating agreement compared to the intersubject consistency of ratings

To estimate the agreement between GPT-4V and human social perceptual ratings in more detail, we calculated the Pearson correlation between the GPT-4V ratings and the average ratings of humans for each social feature which we simply call “agreement of GPT-4V”. To compare to agreement of GPT-4V with the consistency of ratings between different human individuals, we calculated the Pearson correlation of a single human participant’s rating with the average ratings of all others, which we call “intersubject consistency”. Hence, the comparison between agreement of GPT-4V and intersubject consistency estimates the accuracy of social perception of GPT-4V against to that of a randomly chosen human participant.

**Figure 3** (top row) shows the agreement of GPT-4V against the intersubject consistency for image annotations. The average agreement of GPT-4V over all social features was 0.74 and feature specific correlations ranged between 0.41 (Compliant) and 0.95 (Laying down). All feature-specific correlations were statistically significant (p<0.001). The average intersubject consistency was 0.59 and for 95 % of the features (130 / 136) the agreement of GPT-4V was higher than the intersubject consistency. This indicates that for almost all features, GPT-4V derived ratings from images were more reliable population level estimates than subjective evaluations of a single human participant. For Compliant, Physically aggressive, Sexually aroused, Smelling something, Eating or drinking, and Masculine the intersubject consistency was slightly higher than the GPT-4V agreement. Overall, there was strong association between the agreement of GPT-4V and the intersubject consistency - the same features were evaluated with high or low correlation relatively similarly in both data (**Figure 3**).

**Figure 3.**
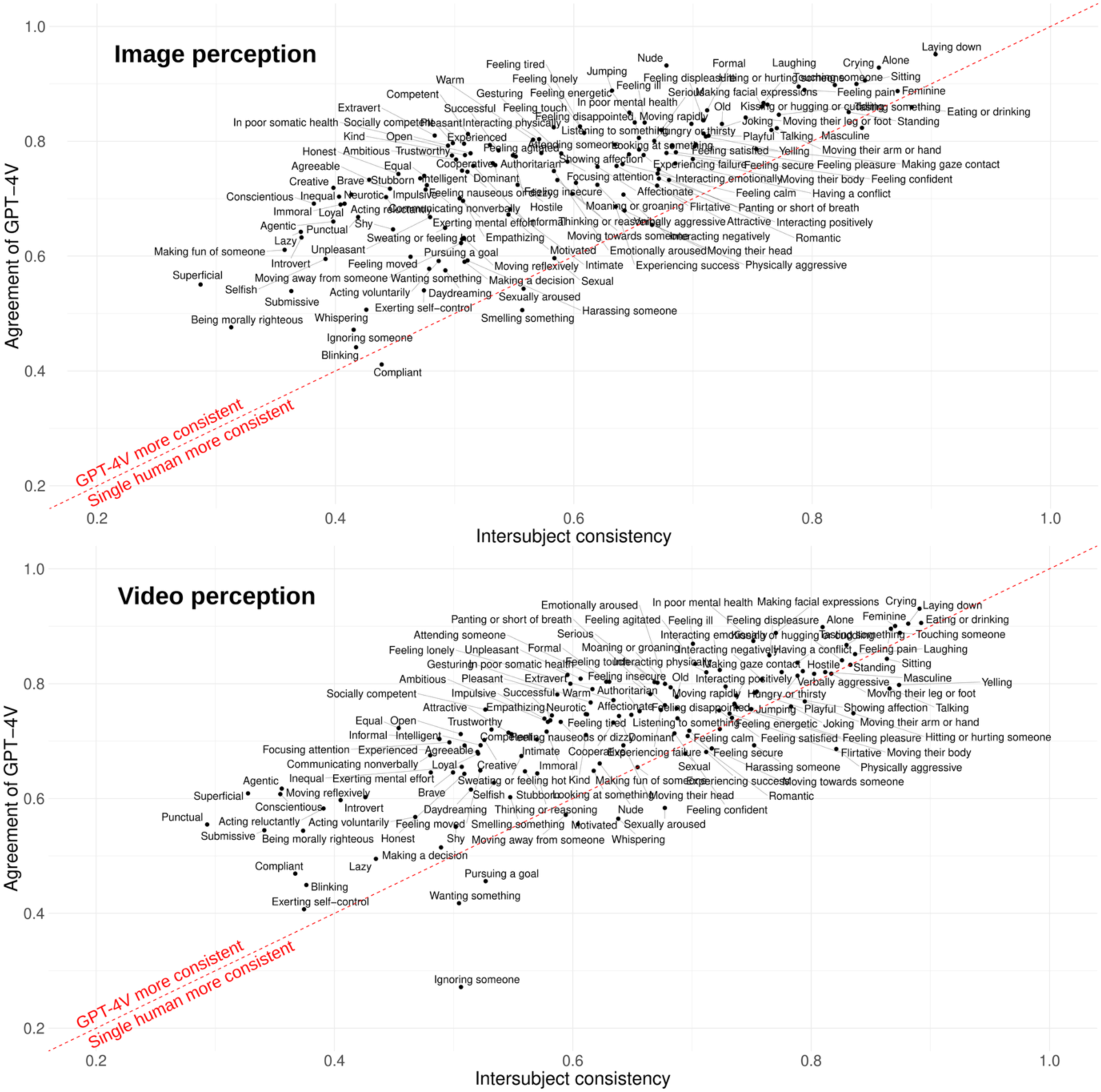
Feature specific rating similarity between GPT-4V and humans for images (top) and videos (bottom). X-axis shows the Pearson correlation of an individual human observers’ ratings (averaged over all individual participants) against the average ratings of all other observers (intersubject consistency), and Y-axis shows the correlation between the GPT-4V ratings and the human average ratings (agreement of GPT-4V). Feature position above the red line indicates that the GPT-4V ratings (averaged over five rounds of data collection) were more consistent estimates of the population level perceptual ratings than those of individual human participants.

**Figure 3** (bottom row) shows the agreement of GPT-4V and the intersubject consistency for the video annotations. The average agreement of GPT-4V over all social features was 0.74 and feature specific correlations ranged between 0.27 (Ignoring someone) and 0.93 (Crying). All feature-specific correlations were statistically significant (p<0.001). The average intersubject consistency was 0.66 and for 85 % of the features (116 / 136) the agreement of GPT-4V was higher than the intersubject consistency. This indicates that for most of the features, GPT-4V derived ratings from videos were more consistent population level estimates than subjective evaluations of a single human participant. For a few features the GPT-V4 rating agreement was considerably lower than the intersubject consistency (Ignoring someone, Wanting something, Yelling, Whispering, Talking, Hitting or hurting someone, Physically aggressive, and Sexually aroused).

Since a single human’s ratings are rarely used as stable population level estimates, we also compared the agreement of GPT-4V with the group-level consistency calculated as the average correlation between any two groups or five human annotations. This group-level consistency for the average rating of five humans was 0.75 (GPT-4V agreement = 0.74) for images and 0.81 (GPT-4V agreement = 0.74) for videos. The agreement of GPT-4V was higher than the group-level consistency for 43 % of the features for images and 17 % for the videos.

### Similarity of the social feature representations between GPT-4V and humans

The social feature annotation structure was consistent between GPT-4V and humans. Feature-wise correlation matrices were structurally similar between GPT-4V and human observer datasets for both images and videos (r_image_ = 0.89, p < 10^−6^, r_video_ = 0.87, p < 10^−6^ based on the Mantel test for similarity matrices, **Figure 4**, left panel). The principal coordinate analysis revealed that mostly similar principal components (PC) emerged from the correlation structure of GPT-4V and human social feature rating data. These similarities are indicated by significant correlations (p < 0.001) between the corresponding PC’s loadings of GPT-4V and human observer data for the first few PCs (**Figure 4**, right panel, clear diagonal for first PCs). The original study with human annotations showed that eight perceptual components emerge from this human data (Santavirta et al., 2024) and rest of the PCs merely model random noise. Hence, it was expected that the link (diagonal correlations) between human and GPT-4V derived components should vanish after around eight PCs which indeed was the case. These results indicate that the structural representation of GPT-4V follows closely the social perceptual representation of humans.

**Figure 4.**
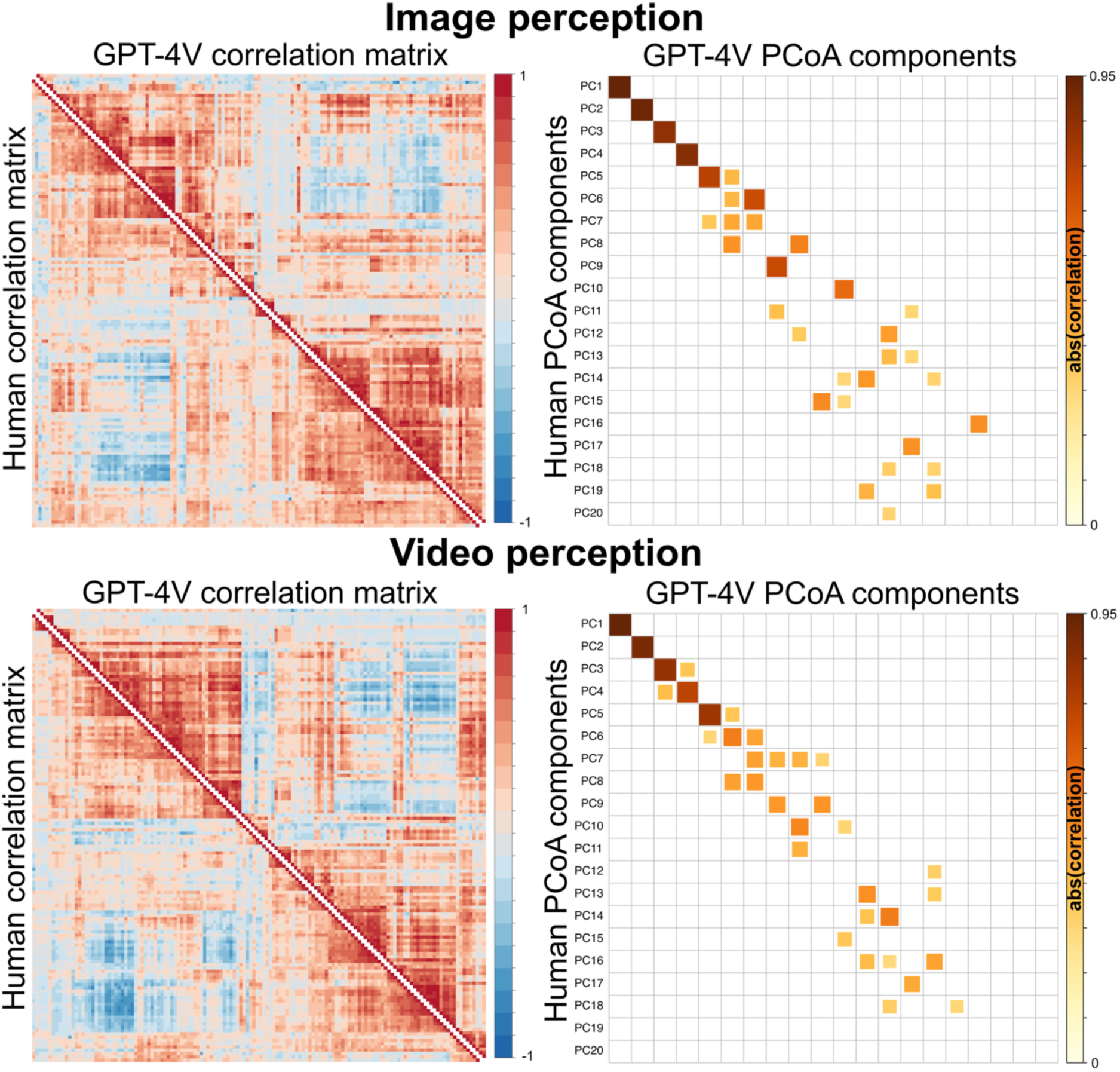
Similarity of the social feature representations for images (top row) and videos (bottom row) between GPT-4V and humans. Leftmost plots show the pairwise correlations of all 136 analyzed social features in the GPT-4V and human rating data. All correlation matrices have been sorted to the same order based on the hierarchically ordered human correlation matrix of the video dataset. Rightmost plots show the similarity of the dimensionality in social perception based on principal coordinate analysis. Significant correlations of the loadings for 20 first principal components between GPT-4V and human derived results are shown (p < 0.001). Absolute correlations are plotted since either high positive or high negative correlation between the component loadings between GPT-4V and humans indicate that the component contains similar information because signs of the components are arbitrary.

### The similarity of the neural representations for social features between GPT-4V and humans

To first evaluate the whole-brain similarity of the response patterns, we calculated the spatial correlations between the unthresholded population level beta coefficient maps between GPT-4V and human based analyses (**Figure 5**, top barplot). The mean correlation across features was 0.92 (range: 0.29 [Nude] – 1.00 [Making facial expressions]) and the correlation was over 0.7 for 96 % of the social features. Next, we calculated the agreement between the statistically thresholded GPT-4V results by calculating the PPVs and NPVs when considering the human derived results as the ground truth. For conservative threshold (voxel-level FWE-corrected, p < 0.05) the mean PPV was 0.73 (range: 0.04 [Feeling lonely] – 1.00 [Old]) and for the lenient threshold (p < 0.001, uncorrected) the mean PPV was 0.74 (range: 0.11 [Feeling lonely] – 1.00 [Old]). The mean NPV for the conservative threshold was 0.97 (range: 0.83 [Moaning] – 1.00 [Feeling tired]) and 0.95 for the lenient threshold (range: 0.77 [Moaning] – 1.00 [Kissing or hugging]). **Figure 5** (bottom barplot) visualizes the feature specific PPVs. See **Figure SI-4**, for brain response patterns of selected social features.

**Figure 5.**
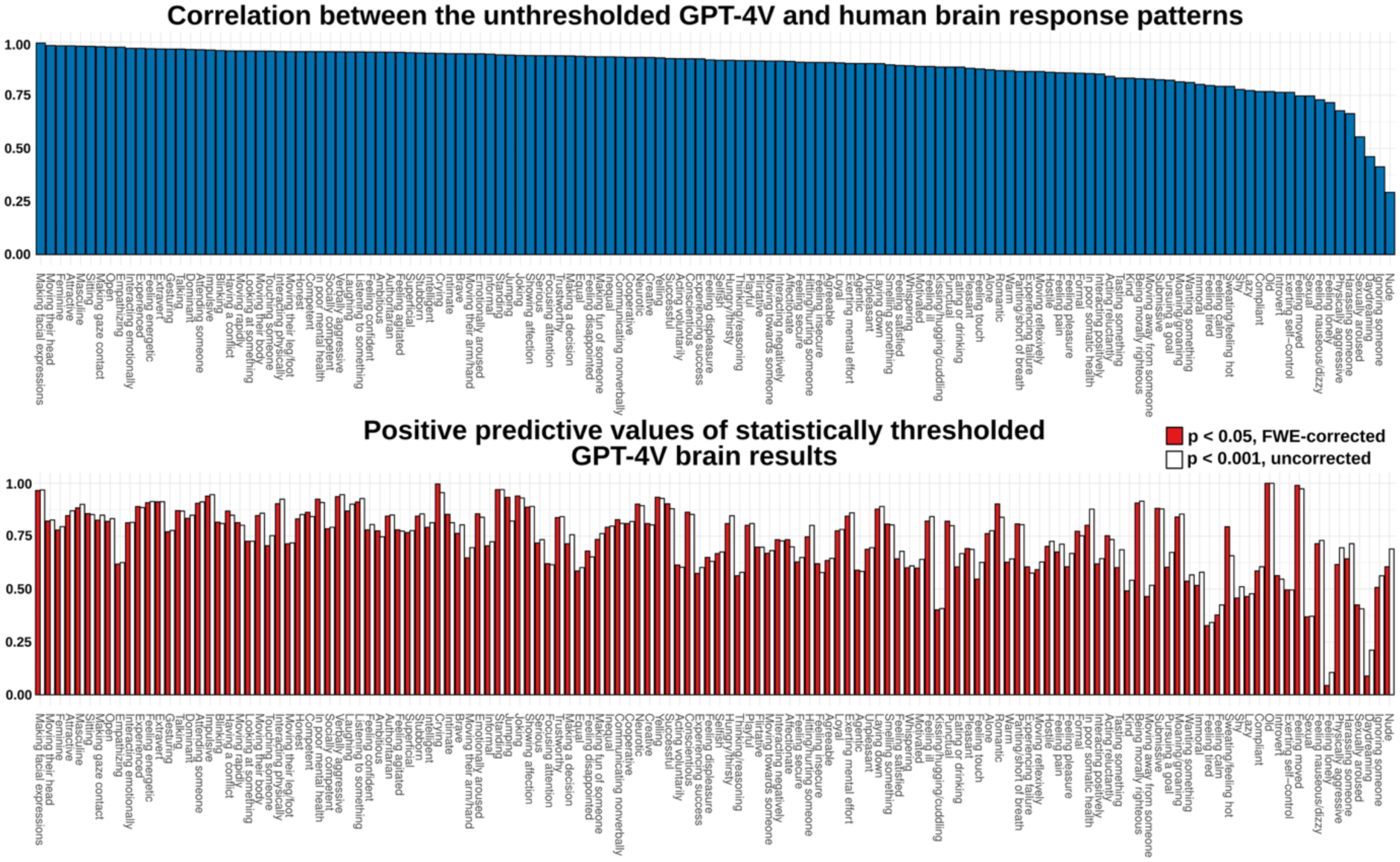
Feature-specific agreement between the brain activation patterns modeled by GPT-4V and human observer responses. Upper plot shows the whole brain response pattern similarity (spatial correlation between the population level unthresholded beta coefficients for each social feature) between GPT-4V and human based analyses. Lower plot shows the statistically thresholded positive predictive values of the GPT-4V results (for the positive association between BOLD signal and social feature). PPVs are plotted for conservative threshold (p < 0.05, voxel-level FWE-corrected) and for lenient threshold (p < 0.001, uncorrected).

### Neural representation for social perception based on GPT-4V stimulus models

**Figure 6** shows the cumulative brain activation maps that were generated by calculating the sum of features that associated positively with hemodynamic response in each voxel (p < 0.001, uncorrected). These maps thus reflect i) whether a given voxel responds to social information at all and ii) whether it’s tuning for social features is narrow (responds to only few features) or wide (responds to multiple features). The cumulative map for GPT-4V was similar with the human derived social cumulative map (r = 0.95) highlighting the general social perceptual network involving lateral occipitotemporal cortex (LOTC), superior temporal sulcus (STS), fusiform gyrus (FG), temporoparietal junction (TPJ), inferior frontal gyrus (IFG) and visual & auditory cortices. While the overall similarity was high, the GPT-4V based cumulative social map showed more associations between social features and hemodynamic response in many areas of the social perceptual network (**Figure 6**, right surface maps).

**Figure 6.**
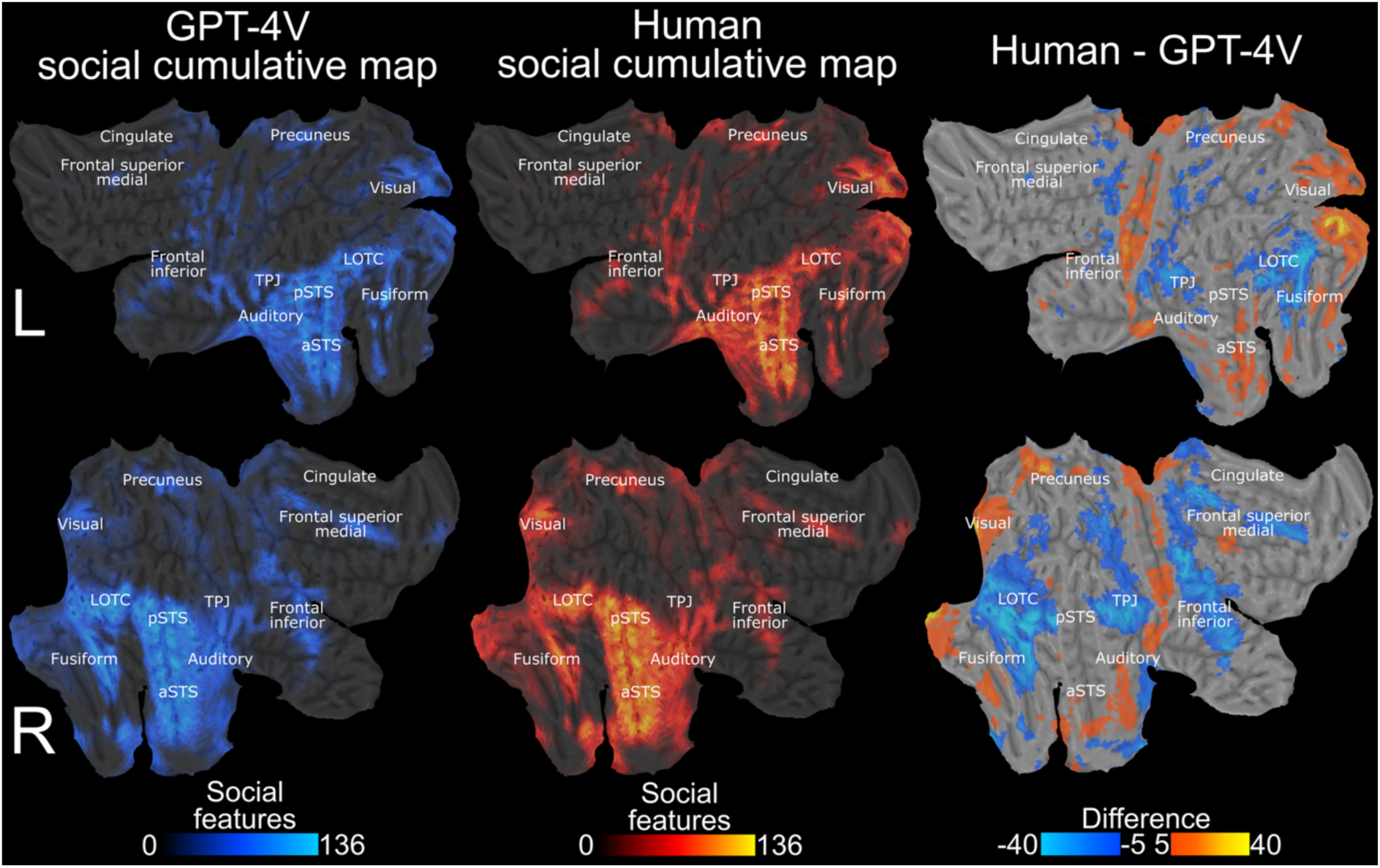
Neural representational space for social perception with stimulus models derived via GPT-4V and human observers. The brain surface maps show how many social features (out of all 136 social features) were associated positively (p < 0.001, uncorrected) with the BOLD response in the given voxel. Leftmost column shows brain maps with GPT-4V-derived stimulus models while middle column shows results with stimulus models based on human observers. Rightmost column shows the absolute difference between cumulative maps highlighting areas where human (hot colours) and GPT-4V (cold colours) models produced a larger number of significant associations with social features. LOTC: Lateral occipitotemporal cortex, pSTS: posterior superior temporal sulcus, aSTS: anterior superior temporal sulcus, TPJ: temporoparietal junction.

## Discussion

Our main findings were that i) GPT-4V is capable of human-like social annotation of complex and dynamic social movie scenes, ii) the annotation capabilities go beyond simple visual social features and, iii) the similarity between human and GPT-4V performance is observed at the perceptual and neural levels. Like humans, GPT-4V evaluated inferred and abstract features with lower agreement with the humans while observable and concrete features were evaluated with high agreement. The structure of the social perceptual space aligned with the human social perceptual space and the similarities of annotations also yielded similar neural representations between GPT-4V and human based stimulus models for dynamic social stimuli. Previous psychological research with AI has mainly focused on text-based tasks for studying the similarity between LLMs and humans (Bodroza et al., 2023; Pan & Zeng, 2023; Strachan et al., 2024) or has transformed behavioral tasks into textual vignettes (Binz & Schulz, 2023; Huang et al., 2023; Tak & Gratch, 2024; X. Wang et al., 2023). To our knowledge, our results are the first evidence of high-level visual perceptual capabilities of LLMs in psychological context. These findings suggest that the current AI models have advanced capabilities to rate the presence of social features that exceed simple feature recognition, suggesting that these models could be used in numerous applications ranging from surveillance to patient monitoring, analysis of customer behavior and social robotics. Our results also provide a framework for how the visual capabilities of GPT-4V can be utilized for studying social cognition both at the perceptual and neural levels, potentially augmenting human annotations or even replacing humans when reliable automated annotations can be achieved (Dillion et al., 2023). Importantly, laborious and costly human experiments could be conducted to test the most promising hypotheses that were first derived with preliminary analyses using LLM annotations.

### Consistent social feature ratings across GPT-4V and human observers

These results were achieved by collecting GPT-4V data five times and by averaging the evaluations over the collection rounds. The overall similarity of the social feature ratings between GPT-4V and human average was high (0.79 for both images and videos). For the low ratings (rating =< 5), GPT-4V gave generally lower values compared to humans, but this bias was not evident in higher ratings (**Figure 2**), indicating that humans evaluate the subtle presence of social features somewhat more effectively than GPT-4V.

The agreement of GPT-4V (the correlation between GPT-4V ratings with the average of humans) was superior compared to the intersubject consistency (the correlation between a single participant’s ratings with the average of others) in social feature evaluations (**Figure 3**). This indicates that the GPT-4V ratings are more accurate estimates of the population average as those of single human observers. GPT-4V evaluations excelled the intersubject consistency for 95 % (images) or 85 % (videos) of the social features.

Although GPT-4V annotations excelled those of single humans, the average annotations over five humans (group-level consistency) still excelled the GPT-4V annotations. However, the difference in reliability was small. The group-level human consistencies were 0.75 (images) and 0.81 (videos), while the matching agreements of GPT-4V were 0.74 in both cases. The agreement of GPT-4V was even higher than the group-level consistency for 43 % (images) and 17 % (videos) of the social features. This indicates that the GPT-4V annotation reliability for images was similar compared to group average calculated from five human participants while the reliability in video annotation was slightly lower.

### Consistent structural representation of social features across GPT-4V and human observers

Both image and video datasets revealed high concordance in the low-level dimensional representations of the social scenes (**Figure 4**). First few orthogonal principal components derived from GPT-4V data in the principal coordinate analysis contained similar information with mainly one corresponding PC in the human results. Of the components, the first eight PCs were previously found to explain meaningful variance in the perceptual data and the rest are likely modeling random noise (Santavirta et al., 2024). As expected, there was thus no clear correspondence between GPT-4V and human PCs in latter components. Based on the detailed examinations of the human data in the original experiment, the eight identified perceptual dimensions were: Unpleasant - Pleasant (PC1), Empathetic - Dominant (PC2), Physical - Cognitive (PC3), Disengaged - Loyal (PC4), Introvert - Extravert (PC5), Playful - Sexual (PC6), Alone - Together (PC7) and Feminine - Masculine (PC8) (Santavirta et al., 2024). The first five PCs showed higher concordance than the rest, indicating that GPT-4V was able to pick the most fundamental components well. This may be due to the fact that the these social perceptual dimensions have long been known to social psychology (Evans & Stanovich, 2013; Goldberg, 1990; Maner, 2017; Oosterhof & Todorov, 2008; Osgood & Suci, 1955; Zajonc, 1980). The similarity of the social annotation structure was also high when measured as the raw correlation between the correlation matrices of the social features ratings (r_image_ = 0.89, p < 10^−6^, r_video_ = 0.87, p < 10^−6^). **Figure 4** (correlation matrices on the left) shows this high similarity visually, while it can also be seen that the human data has some more fine-grained structure compared to the GPT-4V data. These results indicate that GPT-4V annotations follow similar representational structure as human annotations do, while GPT-4V representation lacks some fine-grained details of the human perceptual space (at least in this population of human raters). Importantly, the results emerge from dynamic stimuli where people and faces can be presented in varying positions and orientations instead of e.g. standardized facial expressions commonly used in studies of social perception.

### Neural representations of social perception modeled with GPT-4V and human annotations

To show the usefulness and reliability of GPT-4V annotations for cognitive neuroscience we modeled a retrospective fMRI dataset with stimulations models derived from these annotations. Agreement in social feature annotations also extended to the neural level as was expected from the robust behavioural convergence. The cumulative neural maps, that highlight the general network for social perception in the human brain, were very similar between GPT-4V and humans (r = 0.95, **Figure 6**) highlighting the general social perceptual network involving lateral occipitotemporal cortex (LOTC), superior temporal sulcus (STS), fusiform gyrus (FG), temporoparietal junction (TPJ), inferior frontal gyrus (IFG) and visual & auditory cortices. Previous studies have shown that these regions are involved in social perception (Lahnakoski et al., 2012; Lee Masson & Isik, 2021; McMahon et al., 2023; Nummenmaa & Calder, 2009; Pitcher & Ungerleider, 2021; Santavirta et al., 2023), face perception (Kanwisher & Yovel, 2006; Kessler et al., 2011; Wegrzyn et al., 2015) and language and action perception (Lingnau & Downing, 2015; Wurm & Caramazza, 2019; Wurm & Lingnau, 2015). The cumulative results with GPT-4V stimulus models showed more associations in social perceptual areas LOTC, TPJ, and IFG, while both GPT-4V and humans highlighted the STS region similarly. STS is considered the general hub for abstract social perception (Deen et al., 2015; Lahnakoski et al., 2012; Pitcher & Ungerleider, 2021).

The feature-specific neural representations were also consistent between GPT-4V and humans for most social features (correlation of the unthresholded beta maps = 0.89) but for six social features the correlations were considerably lower than for others (r < 0.70, **Figure 5**, Nude, Sexually aroused, Daydreaming, Ignoring someone, Harassing someone, Physically aggressive). For all these features even a single participant’s ratings would have been more reliable population level estimates as the GPT-4V rating in the video experiment, which most likely explains the weaker consistency in the neural representations of these social features. The statistically thresholded results showed similar levels of consistency than the unthresholded results. On average, the positive predictive value for GPT-4V derived positive associations between social feature and voxelwise brain responses was 0.73 (voxel-level FWE-corrected, p < 0.05) and 0.74 (p < 0.001, uncorrected) and the average negative predictive values were 0.97 and 0.95, respectively. Together these results indicate that the GPT-4V derived social feature ratings from video stimuli were meaningful and produced similar neural representation than human evaluations for most rated social features. These results can be used as a reference for identifying social features that are reliably perceived by GPT-4V.

Because the fMRI responses modeled by GPT-4V and human annotations are consistent, human annotations could be augmented or even replaced in neuroimaging experiments warranting high-dimensional and laborious stimulus mappings. Such automated annotation would allow generation of high-dimensional stimulus models for already collected neuroimaging datasets currently lacking detailed annotations, which would enable efficient mapping of representational spaces with high statistical power (Huth et al., 2016, 2012; Koide-Majima et al., 2020; Lettieri et al., 2019; Saarimäki et al., 2025; Santavirta et al., 2023; Tarhan & Konkle, 2020). This also leads to significant financial advantages: The data collection cost for the 2254 online participants was over $10 000 in participant compensations and required altogether over 1100 hours of labor from the volunteers (∼ 30 min / participant), whereas collecting similar annotations from GPT-4V API cost $100 (< 1 % of the cost). This efficiency allows scaling up the stimulus annotations in neuroimaging experiments significantly.

### Consistency across image and video stimuli

The overall annotation agreement of GPT-4V was equally good for both image and video annotation (r_image_ = 0.79 vs. r_video_ = 0.79). When comparing the feature-specific agreement of GPT-4V with the intersubject consistency (reliability of a single human) or group-level consistency (reliability of five humans) the video annotations were slightly inferior compared to the image annotations. The GPT-4V agreement excelled intersubject consistency for 95% (images) versus 85 % (videos) of the features while GPT-4V agreement was superior to group-level consistency (the average of five humans) for 43 % (images) versus 17 % (videos) of the features. The videos were decomposed into eight frames and a transcript for GPT-4V annotation which limited the available information compared to human subjects. Even with such simplification, the results showed near human-level annotations for concrete and complex social features and robust alignment in the resulting neuroimaging results. The observed inferiority is likely (at least partly) a consequence of simplified input rather than significant inferiority in annotation capability. With the fast development of LLMs, we anticipate that more realistic video input to the LLMs is possible soon and we would expect even better reliability in video annotation when this becomes possible.

Overall, image and video annotation data indicated a similar gradient in both GPT-4V and human data from high agreement in more observable social features (Eating/drinking, Sitting, Talking, Laying down, Touching someone, Laughing, Crying, Making facial expressions) to lower agreement in more complex inferred social features (Conscientious, Superficial, Submissive, Compliant, Acting reluctantly, Being morally righteous) (**Figure 3**). The features with the lowest intersubject consistency may be difficult to infer from static images or short video clips. They may require accumulating information from longer time periods, or their evaluation may be inherently subjective depending on subject-specific properties, such as their previous experiences. Interestingly, the agreement of GPT-4V was relatively higher than the intersubject consistency for the most subjectively evaluated features compared to the features with high intersubject consistency (**Figure 3**). This suggests that although their evaluation is largely subjective, GPT-4V was able to capture the population level inference even for these features.

### The effect of the chosen model

The data were collected using the newest GPT-4.1 model (gpt-4.1-2025-04-14). The image and video annotation data were initially collected using older models (images: gpt-4-1106-vision-preview, videos: GPT-4-turbo-2024-04-09). The summary of the results and comparison with the GPT-4.1 model is shown in **Table SI-3**. On the behavioral level, the GPT-4.1 model achieved higher agreement with humans compared to the older models (images: r_GTP-4_ = 0.71, r_GTP-4.1_ = 0.79, videos: r_GTP-4 Turbo_ = 0.63, r_GTP-4.1_ = 0.79). The brain-level results were consistent across the used models (cumulative maps: r_GTP-4 Turbo_ = 0.94, r_GTP-4.1_ = 0.95). A notable difference was that the GPT-4 Turbo was able to provide ratings for all videos while the GPT-4.1 refused to provide ratings for scenes containing sexual material. In the image experiment, both used models failed to rate some images containing sex and blood, indicative of internal filters for material that some users might find objectionable. While the GPT-4.1 model showed overall improvement of the results, the content moderation for sexual content was also improved limiting the use cases of the model.

We used closed-source GPT-models, which are all likely to be replace with new models in the future. As new models are developed, it is unlikely that the observed capabilities would disappear as we see improvement with the new GPT 4.1 model compared to the previous one. We would also expect to observe similar annotation capabilities with other developers’ flagship models. A bigger concern would be the deliberate moderation of usage and accepted input content, and an ideal solution would involve an open-source LLM without any content filters. However, locally installed LLMs would require more computational resources than are currently available for most users, even in research. For these reasons, closed-source models such as GPT-4V offer a good compromise for now.

### Future directions

First, because human observers and GPT-4V provided inconsistent responses for some features, future studies need to quantify the sources of variation that would explain the differences between GPT-4V and human perceptual ratings. One possible way to increase the reliability of the LLM derived results could be to harmonize the perceptual data collection by assigning different sociodemographic or personality characteristics to the LLM (Argyle et al., 2023) before collecting perceptual evaluations. This could yield even more reliable results in specific subpopulations and perhaps also for population level averages. By manual balancing of the studied subpopulations the results could generalize better to underrepresented groups in the LLM’s training data. Second, these findings show the potential of LLM vision in complex psychosocial phenomena and thus open new avenues for studying the LLM perception. Future research should investigate whether LLMs are able to reliably infer, for example, emotions or personalities just by perceiving them in dynamic settings. If LLM perception proves to be reliable, it could be used in countless ways to simulate experiments, which would free resources to confirm findings with human subjects only in the most potential study designs (Dillion et al., 2023; Horton, 2023). Ultimately, LLM perception could, for example, help medical professionals to identify the patients in acute distress or LLMs could monitor people in real-life to give physicians more insights of people’s psychological and physical health by their behavior in everyday situations. Previous machine learning models are trained for very specific purposes and their use require expertise. The potential in LLMs relies on the fact that they are accessible to wide audiences and can be used for many purposes without additional training and with little cost.

### Limitations

We compared the agreement of GPT-4V social evaluations against the agreement of single human participants and groups of five participants, yet usually more human participants are needed for accurate population level estimates. However, this approach allows answering the question of whether GPT-4V performs at a level comparable to a single or few subjects randomly drawn from the sample, and the answer to this question is “yes”. Our data with ten human ratings for each item is not large enough to enable random sampling of larger subsets of human annotations. Importantly, the GPT-based stimulus model yielded comparable neural representations of social features as that based on humans, indicating population-level accuracy of the model.

The specific details of the input prompt may influence the LLM’s output considerably (L. Wang et al., 2024). We cannot estimate how much the results would change and how with differing prompts. We designed the prompt to be as close to the human instructions as possible while at the same time ensuring that GPT-4V provides evaluations in structured format. In a pilot study, we showed that minor changes in the prompt did not have major influence on the evaluations (see section “The stability of GPT-4V outputs with slightly different prompts” in Supplemental Materials).

LLM outputs are subject to the underlying biases in the training data. Therefore, the LLM’s output may only represent a specific sample of the population, and it cannot capture the opinion that is not represented in the training data (Demszky et al., 2023). For example, the unguided opinions of many LLMs are more inclined to reflect the opinions of individuals who are liberal, high income, well-educated and not religious in one study (Santurkar et al., 2023) and the opinions were interpreted as right-leaning in another (Park et al., 2024) while another study with GPT-4 model described the political views as moderate (Almeida et al., 2024). Nevertheless, LLMs can produce accurate opinions of different sociodemographic groups when guided to do so (Argyle et al., 2023) but seems to have tendency to favor one’s own group (ingroup solidarity) and derogate other groups (outgroup hostility) (Hu et al., 2024). It is not known whether these biases majorly influence the current findings, since this study focuses on mainly observable and instant perceptions of social interactions, not political opinions that can differ drastically between people. Previous study of a taxonomy for social perception in humans could not find evidence that the sex, age or ethnicity of the perceiver would have major effect on the perceptual evaluations of the current movie clip stimulus (Santavirta et al., 2024).

It is likely that content associated with mixed moral opinions is deliberately left out from the training data of commercially available LLMs and the models also incorporate additional content filters to prevent harmful use (OpenAI et al., 2023). The GPT-4 model documentation reports several content filters for the input including “sexual”, “hate”, “harassment”, and “violence” filters (https://platform.openai.com/docs/api-reference/moderations/create). Due to these content filters, we could not collect data for most images containing sex (and for two images containing blood). The outputs of the GPT-4 are also adjusted, for example, not to encourage people to commit crimes (OpenAI et al., 2023). These output moderations may have influenced the perceptual evaluations of some disturbing scenes. Content filtering most likely explains why the neural representations for features “Harassing someone”, “Physically aggressive” and “Sexually aroused” had the weakest correspondence of all features with human representations.

Finally, the results do not reveal how GPT-4V, or humans, make the judgments and give their final evaluations for the presence of social features in the images or videos of social interactions. Although the evaluations between GPT-4V and human align well, it does not indicate that GPT-4V would perceive or understand the situations similarly as humans. The results simply indicate that GPT-4V can be used as a practical tool for collecting human-like annotations of even complex social features. The same is true for the alignment of the modelled neural responses. The results do not indicate that GPT-4V would process information similar to these brain regions in the human brain. Instead, it indicates that GPT-4V based models can be used to create predictive models for naturalistic neuroimaging data.

## Conclusion

Our work provides first evidence of the advanced social feature annotations of GPT-4V from images and videos. The structural representation of GPT-4V for social features aligned with the representations of humans and the GPT-4V based stimulus models also yielded in similar neural representations compared to gold-standard human based models. The general human social perceptual network including mainly occipitotemporal brain regions was clearly identified by using stimulus model based on GPT-4V annotations. The results suggest that in the future, AI can be used for supporting a multitude of applications requiring complex visual inference ranging from patient monitoring to surveillance and customer behavior tracking. Our results also suggest that LLMs can be used for generating rich stimulus models for high-dimensional representational mapping of brain functions with fMRI, also allowing large-scale reanalysis of existing fMRI datasets where high-dimensional stimulus models can be retrieved post hoc using LLMs. However, caution is still warranted generalizing LLM derived results of higher mental processes to reality and future research should experiment with LLM in other domains of social cognition, such as emotion and motivation.

## Data and code availability

The anonymized GPT-4V and human rating data and the data collection and analysis scripts are available in the project’s GitHub repository (https://github.com/santavis/GPT-4V-social-perception). According to Finnish legislation, the original (even anonymized) neuroimaging data used in the experiment cannot be released for public use. The voxelwise (unthresholded) result maps from fMRI analyses can be requested from the authors. The stimulus movie clips can be made available for researchers upon request, but copyrights preclude public redistribution of the stimulus set. Short descriptions of each movie clip can be found in the supplementary materials (**Table SI-1**).

## CRediT authorship contribution statement

**Severi Santavirta**: Conceptualization, Methodology, Software, Validation, Formal analysis, Investigation, Resources, Data curation, Writing – original draft, Writing – review & editing, Visualization, Project administration. **Yuhang Wu**: Conceptualization, Methodology, Software, Validation, Formal analysis, Investigation, Resources, Data curation, Writing – original draft, Writing – review & editing, Visualization. **Lauri Suominen**: Methodology, Software, Investigation, Data curation, Writing – review & editing. **Lauri Nummenmaa**: Conceptualization, Methodology, Resources, Writing – original draft, Writing – review & editing, Visualization, Supervision, Project administration.

## Acknowledgements

The study was supported by Turku University Foundation and Alfred Kordelin Foundation grants to SS and Finnish Governmental Research Funding for Turku University Hospital and for the Western Finland collaborative area to SS. We would like to express our gratitude to Haoming Zhong, a graduate student at National Key Laboratory for Novel Software Technology, Nanjing University for his valuable insights, inspiration, and programming assistance on this project.

## References

Adolphs, R., Nummenmaa, L., Todorov, A., & Haxby, J. V. (2016). Data-driven approaches in the investigation of social perception. Philosophical Transactions of the Royal Society of London. Series B, Biological Sciences, 371(1693). 10.1098/rstb.2015.0367

Aher, G. V., Arriaga, R. I., & Kalai, A. T. (2023). Using Large Language Models to Simulate Multiple Humans and Replicate Human Subject Studies. In A. Krause, E. Brunskill, K. Cho, B. Engelhardt, S. Sabato, & J. Scarlett (Eds.), Proceedings of the 40th International Conference on Machine Learning (Vol. 202, pp. 337–371). PMLR. https://proceedings.mlr.press/v202/aher23a.html

Almeida, G. F. C. F., Nunes, J. L., Engelmann, N., Wiegmann, A., & Araújo, M. de. (2024). Exploring the psychology of LLMs’ moral and legal reasoning. Artificial Intelligence, 333, 104145. 10.1016/j.artint.2024.104145

Argyle, L. P., Busby, E. C., Fulda, N., Gubler, J. R., Rytting, C., & Wingate, D. (2023). Out of One, Many: Using Language Models to Simulate Human Samples. Political Analysis: An Annual Publication of the Methodology Section of the American Political Science Association, 31(3), 337–351. 10.1017/pan.2023.2

Binz, M., & Schulz, E. (2023). Using cognitive psychology to understand GPT-3. Proceedings of the National Academy of Sciences of the United States of America, 120(6), e2218523120. 10.1073/pnas.2218523120

Bodroza, B., Dinic, B. M., & Bojic, L. (2023). Personality testing of GPT-3: Limited temporal reliability, but highlighted social desirability of GPT-3’s personality instruments results. In arXiv [cs.AI]. arXiv. http://arxiv.org/abs/2306.04308

Bubeck, S., Chadrasekaran, V., Eldan, R., Gehrke, J., Horvitz, E., Kamar, E., Lee, P., Lee, Y. T., Li, Y., Lundberg, S., Nori, H., Palangi, H., Ribeiro, M. T., & Zhang, Y. (2023). Sparks of Artificial General Intelligence: Early Experiments with GPT-4. ArXiv. 10.48550/arXiv.2303.12712

Deen, B., Koldewyn, K., Kanwisher, N., & Saxe, R. (2015). Functional Organization of Social Perception and Cognition in the Superior Temporal Sulcus. Cerebral Cortex, 25(11), 4596–4609. 10.1093/cercor/bhv111

Demszky, D., Yang, D., Yeager, D. S., Bryan, C. J., Clapper, M., Chandhok, S., Eichstaedt, J. C., Hecht, C., Jamieson, J., Johnson, M., Jones, M., Krettek-Cobb, D., Lai, L., JonesMitchell, N., Ong, D. C., Dweck, C. S., Gross, J. J., & Pennebaker, J. W. (2023). Using large language models in psychology. Nature Reviews Psychology, 2(11), 688–701. 10.1038/s44159-023-00241-5

Dillion, D., Tandon, N., Gu, Y., & Gray, K. (2023). Can AI language models replace human participants? Trends in Cognitive Sciences, 27(7), 597–600. 10.1016/j.tics.2023.04.008

Esteban, O., Markiewicz, C. J., Blair, R. W., Moodie, C. A., Isik, A. I., Erramuzpe, A., Kent, J. D., Goncalves, M., DuPre, E., Snyder, M., Oya, H., Ghosh, S. S., Wright, J., Durnez, J., Poldrack, R. A., & Gorgolewski, K. J. (2019). fMRIPrep: a robust preprocessing pipeline for functional MRI. Nature Methods, 16(1), 111–116. 10.1038/s41592-018-0235-4

Evans, J. S. B. T., & Stanovich, K. E. (2013). Dual-Process Theories of Higher Cognition: Advancing the Debate. Perspectives on Psychological Science: A Journal of the Association for Psychological Science, 8(3), 223–241. 10.1177/1745691612460685

Fonov, V. S., Evans, A. C., McKinstry, R. C., Almli, C. R., & Collins, D. L. (2009). Unbiased nonlinear average age-appropriate brain templates from birth to adulthood. NeuroImage, 47, S102. 10.1016/S1053-8119(09)70884-5

Gemini Team, Reid, M., Savinov, N., Teplyashin, D., Dmitry, Lepikhin, Lillicrap, T., Alayrac, J.-B., Soricut, R., Lazaridou, A., Firat, O., Schrittwieser, J., Antonoglou, I., Anil, R., Borgeaud, S., Dai, A., Millican, K., Dyer, E., Glaese, M., … Vinyals, O. (2024). Gemini 1.5: Unlocking multimodal understanding across millions of tokens of context. In arXiv [cs.CL]. arXiv. http://arxiv.org/abs/2403.05530

Goldberg, L. R. (1990). An alternative “description of personality”: The Big-Five factor structure. Journal of Personality and Social Psychology, 59(6), 1216–1229. 10.1037/0022-3514.59.6.1216

Horton, J. J. (2023). Large Language Models as Simulated Economic Agents: What Can We Learn from Homo Silicus? ArXiv, 31122. 10.48550/arXiv.2301.07543

Hu, T., Kyrychenko, Y., Rathje, S., Collier, N., van der Linden, S., & Roozenbeek, J. (2024). Generative language models exhibit social identity biases. Nature Computational Science. 10.1038/s43588-024-00741-1

Huang, J.-T., Lam, M. H., Li, E. J., Ren, S., Wang, W., Jiao, W., Tu, Z., & Lyu, M. R. (2023). Emotionally Numb or Empathetic? Evaluating How LLMs Feel Using EmotionBench. In arXiv. arXiv. 10.48550/arXiv.2308.03656

Huth, A. G., de Heer, W. A., Griffiths, T. L., Theunissen, F. E., & Gallant, J. L. (2016). Natural speech reveals the semantic maps that tile human cerebral cortex. Nature, 532(7600), 453–458. 10.1038/nature17637

Huth, A. G., Nishimoto, S., Vu, A. T., & Gallant, J. L. (2012). A continuous semantic space describes the representation of thousands of object and action categories across the human brain. Neuron, 76(6), 1210–1224. 10.1016/j.neuron.2012.10.014

Kanwisher, N., & Yovel, G. (2006). The fusiform face area: a cortical region specialized for the perception of faces. Philosophical Transactions of the Royal Society of London. Series B, Biological Sciences, 361(1476), 2109–2128. 10.1098/rstb.2006.1934

Karjalainen, T., Karlsson, H. K., Lahnakoski, J. M., Glerean, E., Nuutila, P., Jaaskelainen, I. P., Hari, R., Sams, M., & Nummenmaa, L. (2017). Dissociable Roles of Cerebral mu-Opioid and Type 2 Dopamine Receptors in Vicarious Pain: A Combined PET-fMRI Study. Cerebral Cortex, 27(8), 4257–4266. 10.1093/cercor/bhx129

Katz, D. M., Bommarito, M. J., Gao, S., & Arredondo, P. (2024). GPT-4 passes the bar exam. Philosophical Transactions. Series A, Mathematical, Physical, and Engineering Sciences, 382(2270), 20230254. 10.1098/rsta.2023.0254

Kessler, H., Doyen-Waldecker, C., Hofer, C., Hoffmann, H., Traue, H. C., & Abler, B. (2011). Neural correlates of the perception of dynamic versus static facial expressions of emotion. Psycho-Social Medicine, 8, Doc03. 10.3205/psm000072

King, M. (2023). Administration of the text-based portions of a general IQ test to five different large language models. TechRxiv. 10.36227/techrxiv.22645561.v1

Koide-Majima, N., Nakai, T., & Nishimoto, S. (2020). Distinct dimensions of emotion in the human brain and their representation on the cortical surface. NeuroImage, 222, 117258. 10.1016/j.neuroimage.2020.117258

Kung, T. H., Cheatham, M., Medenilla, A., Sillos, C., De Leon, L., Elepaño, C., Madriaga, M., Aggabao, R., Diaz-Candido, G., Maningo, J., & Tseng, V. (2023). Performance of ChatGPT on USMLE: Potential for AI-assisted medical education using large language models. PLOS Digital Health, 2(2), e0000198. 10.1371/journal.pdig.0000198

Lahnakoski, J. M., Glerean, E., Salmi, J., Jaaskelainen, I. P., Sams, M., Hari, R., & Nummenmaa, L. (2012). Naturalistic FMRI mapping reveals superior temporal sulcus as the hub for the distributed brain network for social perception. Frontiers in Human Neuroscience, 6, 233. 10.3389/fnhum.2012.00233

Lee Masson, H., & Isik, L. (2021). Functional selectivity for social interaction perception in the human superior temporal sulcus during natural viewing. NeuroImage, 245, 118741. 10.1016/j.neuroimage.2021.118741

Lettieri, G., Handjaras, G., Ricciardi, E., Leo, A., Papale, P., Betta, M., Pietrini, P., & Cecchetti, L. (2019). Emotionotopy in the human right temporo-parietal cortex. Nature Communications, 10(1), 5568. 10.1038/s41467-019-13599-z

Lingnau, A., & Downing, P. E. (2015). The lateral occipitotemporal cortex in action. Trends in Cognitive Sciences, 19(5), 268–277. 10.1016/j.tics.2015.03.006

Maner, J. K. (2017). Dominance and Prestige: A Tale of Two Hierarchies. Current Directions in Psychological Science, 26(6), 526–531. 10.1177/0963721417714323

Mantel, N. (1967). The detection of disease clustering and a generalized regression approach. Cancer Research, 27(2), 209–220. https://www.ncbi.nlm.nih.gov/pubmed/6018555

McMahon, E., Bonner, M. F., & Isik, L. (2023). Hierarchical organization of social action features along the lateral visual pathway. Current Biology: CB, 33(23), 5035–5047.e8. 10.1016/j.cub.2023.10.015

Nummenmaa, L., & Calder, A. J. (2009). Neural mechanisms of social attention. Trends in Cognitive Sciences, 13(3), 135–143. 10.1016/j.tics.2008.12.006

Nummenmaa, L., Lukkarinen, L., Sun, L., Putkinen, V., Seppala, K., Karjalainen, T., Karlsson, H. K., Hudson, M., Venetjoki, N., Salomaa, M., Rautio, P., Hirvonen, J., Lauerma, H., & Tiihonen, J. (2021). Brain Basis of Psychopathy in Criminal Offenders and General Population. Cerebral Cortex, 31(9), 4104–4114. 10.1093/cercor/bhab072

Nummenmaa, L., Malèn, T., Nazari-Farsani, S., Seppälä, K., Sun, L., Santavirta, S., Karlsson, H. K., Hudson, M., Hirvonen, J., Sams, M., Scott, S., & Putkinen, V. (2023). Decoding brain basis of laughter and crying in natural scenes. NeuroImage, 273, 120082. 10.1016/j.neuroimage.2023.120082

Oosterhof, N. N., & Todorov, A. (2008). The functional basis of face evaluation. Proceedings of the National Academy of Sciences of the United States of America, 105(32), 11087–11092. 10.1073/pnas.0805664105

OpenAI, Achiam, J., Adler, S., Agarwal, S., Ahmad, L., Akkaya, I., Aleman, F. L., Almeida, D., Altenschmidt, J., Altman, S., Anadkat, S., Avila, R., Babuschkin, I., Balaji, S., Balcom, V., Baltescu, P., Bao, H., Bavarian, M., Belgum, J., … Zoph, B. (2023). GPT-4 Technical Report. ArXiv. 10.48550/arXiv.2303.08774

Osgood, C. E., & Suci, G. J. (1955). Factor analysis of meaning. Journal of Experimental Psychology, 50(5), 325–338. 10.1037/h0043965

Pan, K., & Zeng, Y. (2023). Do LLMs Possess a Personality? Making the MBTI Test an Amazing Evaluation for Large Language Models. ArXiv. 10.48550/arXiv.2307.16180

Park, P. S., Schoenegger, P., & Zhu, C. (2024). Diminished diversity-of-thought in a standard large language model. Behavior Research Methods. 10.3758/s13428-023-02307-x

Pitcher, D., & Ungerleider, L. G. (2021). Evidence for a Third Visual Pathway Specialized for Social Perception. Trends in Cognitive Sciences, 25(2), 100–110. 10.1016/j.tics.2020.11.006

Pruim, R. H. R., Mennes, M., van Rooij, D., Llera, A., Buitelaar, J. K., & Beckmann, C. F. (2015). ICA-AROMA: A robust ICA-based strategy for removing motion artifacts from fMRI data. NeuroImage, 112, 267–277. 10.1016/j.neuroimage.2015.02.064

Saarimäki, H. (2021). Naturalistic Stimuli in Affective Neuroimaging: A Review. Frontiers in Human Neuroscience, 15, 675068. 10.3389/fnhum.2021.675068

Saarimäki, H., Nummenmaa, L., Volynets, S., Santavirta, S., Aksiuto, A., Sams, M., Jääskeläinen, I. P., & Lahnakoski, J. M. (2025). Cerebral topographies of perceived and felt emotions. Imaging Neuroscience, 3. 10.1162/imag_a_00517

Santavirta, S., Karjalainen, T., Nazari-Farsani, S., Hudson, M., Putkinen, V., Seppälä, K., Sun, L., Glerean, E., Hirvonen, J., Karlsson, H. K., & Nummenmaa, L. (2023). Functional organization of social perception in the human brain. NeuroImage, 120025. 10.1016/j.neuroimage.2023.120025

Santavirta, S., Malén, T., Erdemli, A., & Nummenmaa, L. (2024). A taxonomy for human social perception: Data-driven modeling with cinematic stimuli. Journal of Personality and Social Psychology. 10.1037/pspa0000415

Santurkar, S., Durmus, E., Ladhak, F., Lee, C., Liang, P., & Hashimoto, T. (2023). Whose opinions do language models reflect? International Conference on Machine Learning, abs/2303.17548. 10.48550/arXiv.2303.17548

Sohail, A., & Zhang, L. (2025). Using large language models to facilitate academic work in the psychological sciences. Current Psychology (New Brunswick, N.J.). 10.1007/s12144-025-07438-2

Strachan, J. W. A., Albergo, D., Borghini, G., Pansardi, O., Scaliti, E., Gupta, S., Saxena, K., Rufo, A., Panzeri, S., Manzi, G., Graziano, M. S. A., & Becchio, C. (2024). Testing theory of mind in large language models and humans. Nature Human Behaviour, 8(7), 1285–1295. 10.1038/s41562-024-01882-z

Tak, A. N., & Gratch, J. (2024). GPT-4 emulates average-human emotional cognition from a third-person perspective. In arXiv [cs.AI]. arXiv. http://arxiv.org/abs/2408.13718

Tarhan, L., & Konkle, T. (2020). Sociality and interaction envelope organize visual action representations. Nature Communications, 11(1), 3002. 10.1038/s41467-020-16846-w

Wang, L., Chen, X., Deng, X., Wen, H., You, M., Liu, W., Li, Q., & Li, J. (2024). Prompt engineering in consistency and reliability with the evidence-based guideline for LLMs. NPJ Digital Medicine, 7(1), 41. 10.1038/s41746-024-01029-4

Wang, X., Li, X., Yin, Z., Wu, Y., & Liu, J. (2023). Emotional intelligence of Large Language Models. Journal of Pacific Rim Psychology, 17. 10.1177/18344909231213958

Wang, Y., Zhao, J., Ones, D. S., He, L., & Xu, X. (2025). Evaluating the ability of large language models to emulate personality. Scientific Reports, 15(1), 519. 10.1038/s41598-024-84109-5

Wegrzyn, M., Riehle, M., Labudda, K., Woermann, F., Baumgartner, F., Pollmann, S., Bien, C. G., & Kissler, J. (2015). Investigating the brain basis of facial expression perception using multi-voxel pattern analysis. Cortex; a Journal Devoted to the Study of the Nervous System and Behavior, 69, 131–140. 10.1016/j.cortex.2015.05.003

Wurm, M. F., & Caramazza, A. (2019). Lateral occipitotemporal cortex encodes perceptual components of social actions rather than abstract representations of sociality. NeuroImage, 202, 116153. 10.1016/j.neuroimage.2019.116153

Wurm, M. F., & Lingnau, A. (2015). Decoding actions at different levels of abstraction. The Journal of Neuroscience: The Official Journal of the Society for Neuroscience, 35(20), 7727–7735. 10.1523/JNEUROSCI.0188-15.2015

Yang, Z., Li, L., Lin, K., Wang, J., Lin, C.-C., Liu, Z., & Wang, L. (2023). The Dawn of LMMs: Preliminary Explorations with GPT-4V(ision). ArXiv. 10.48550/arXiv.2309.1742

Zajonc, R. B. (1980). Feeling and thinking: Preferences need no inferences. The American Psychologist, 35(2), 151–175. 10.1037/0003-066X.35.2.151

Ziems, C., Held, W. B., Shaikh, O., Chen, J., Zhang, Z., & Yang, D. (2023). Can large language models transform computational social science? International Conference on Computational Logic, 50, 237–291. 10.1162/coli_a_00502

